# Senataxin is required for optimal class switch recombination to the immunoglobulin A isotype

**DOI:** 10.1101/2025.09.23.678005

**Authors:** Vithushan Surendran, Jonathan Mapletoft, Peter Zeng, Julia Avolio, Sam Afkhami, Joshua J. C. McGrath, Andrew T. Chen, Art Marzok, Jann C. Ang, Kyle Amaral, Yona Tugg, Martin R. Stampfli, Matthew S. Miller

## Abstract

Senataxin (SETX) is an RNA/DNA helicase that plays a pivotal role in transcription, R-loop resolution, and the DNA damage response (DDR). Dysfunction of SETX, however, has been implicated in neurodegenerative diseases, including amyotrophic lateral sclerosis and ataxia with oculomotor apraxia type 2. Importantly, both R-loop resolution and the DDR are essential for class switch recombination (CSR) in B cells, a critical step in generating antibody diversity. Here we demonstrate that *ex vivo* polyclonal stimulation of SETX^+/+^ and SETX^-/-^ mouse splenocytes results in equivalent amounts of IgM- and IgG-secreting B cells but significantly fewer IgA-producing B cells in SETX^-/-^ mice. Additionally, SETX^-/-^ mice generate significantly reduced antigen specific IgA titers following infection with influenza A virus. Sequencing of the IgA-secreting B cell repertoire revealed reduced clonal diversity in SETX^-/-^ mice. SETX^-/-^ mice also had increased R-loop formation within the IgA locus. Collectively, these data highlight an important role for SETX in mediating CSR to IgA. As such, a further understanding of SETX’s role in the immune response is important for expanding our knowledge of both the general immunobiology relating to CSR as well as the immunological phenotypes associated with neurodegenerative diseases associate with mutations in SETX.

**One sentence summary:** A deficiency in senataxin results in impaired class switching to IgA *in vitro* with decreased IgA response to infection and IgA repertoire diversity *in vivo*.

## Introduction

Autosomal recessive cerebellar ataxias (ARCA) are a rare group of neurological disorders characterized by cerebellar atrophy that have no cure (1). One such ARCA, ataxia with oculomotor apraxia type 2 (AOA2), is caused by mutations in senataxin (SETX) and is associated with defects in DNA damage repair (2–4). DNA repair pathways are play essential roles in the generation of immune diversity, including through the process of class switching to different immunoglobulin isotypes in B-cells (5–7). Indeed, in addition to neurological manifestations, patients with ataxia telangiectasia (A-T) – another ARCA associated with DNA repair defects – commonly present with selective IgA deficiency (SIgAD) (8). Given the DNA repair defect associated with AOA2, it is possible that there is an associated cryptic immunodeficiency that has not yet been discovered due to the rarity of the disease (1).

Immunoglobulin class switching is a critical mechanism for generating a diverse immune repertoire. Importantly, class switching is driven by class switch recombination (CSR), a complex process through which B cells irreversibly alter their DNA within the immunoglobulin (Ig) heavy chain (IgH) locus to allow for the switching of antibody isotype (5–7). This recombination event results in the removal of the default constant heavy chain (Cμ [IgM]) and facilitates switching to one of the downstream heavy chains, Cγ (IgG), Cε (IgE), or Cα (IgA) – thereby changing the antibody isotype (5–7). Upstream of the various constant regions are switch (S) regions, and upon initiation of CSR, the S regions are transcribed resulting in the formation of germline transcripts (GLT) (7). GLT formation is dependent upon the cytokines in the B cell microenvironment, which promote the targeting of specific antibody isotype S regions via transcription factor activation (9–11). Key features of S regions include high guanosine (G) content and an increased propensity to generate R-loops (12, 13). When R-loops form within the S regions, single-strand DNA (ssDNA) substrates become available for modification by activation induced cytidine deaminase (AID), which converts cytosine (C) to uracil (U) within DNA (14–17). The resulting nucleotide mismatch, U:G, is processed into double stranded breaks (DSBs) within the S regions (18–21). Ultimately, two distinct S regions are ligated together via classical non-homologous end joining (c-NHEJ), resulting in the production of a new antibody isotype (5–7). Interestingly, a recent study exploring the role of the RNA exosome and RNA helicases, including Mtr4 and SETX, on DNA mutagenesis also reported that the CRISPR-mediated knockout of *SETX* in CH12 cells, a murine B cell line, induced a modest defect in the ability of these cells to become IgA^+^ implicating *SETX* in CSR (22). However, whether this defect occurs in primary B cells, and under physiological conditions remains unclear.

Previous studies have illustrated that the function of SETX is pleiotropic (23–27). The primary role of SETX is to resolve DNA:RNA hybrids, known as R-loops, through its RNA:DNA helicase activity in order to prevent DNA damage and preserve genomic stability (26, 28). SETX has also been demonstrated to suppress the anti-viral transcriptional response following viral infection, probably through its role in R-loop regulation(29). SETX deficiency results in an amplified expression of anti-viral mediators and pro-inflammatory cytokines/chemokines following influenza A virus (IAV) infection (29). However, SETX^-/-^ mice do not spontaneously develop any neurological clinical signs, contrary to what is observed in AOA2 patients (30). Rather, deficiencies in SETX result in infertility of male mice, as a lack of R-loop resolution, and subsequently elevated DNA damage, leads to apoptosis in the spermatocytes – likely through the induction of DSBs within the spermatocyte genome (30). Additionally, SETX has been shown to colocalize with a multitude of proteins involved in the DNA damage response (DDR) (30–34). However, the specific role of SETX in the process of the DDR is not completely clear. As such, we hypothesize that the functions of SETX related to its DNA/RNA helicase activity and regulation of R-loops are essential for CSR.

Herein, we demonstrate that SETX plays an important role in facilitating CSR to IgA. *Ex vivo* polyclonal stimulation of SETX^-/-^ B-cells resulted in a significant decrease in IgA^+^ B-cells five days post-stimulation. A significant reduction in IgA repertoire diversity was observed following vaccination in SETX^-/-^ mice when compared to wildtype controls. Furthermore, a deficiency in SETX resulted in an increase in accumulation of R-loops in the immunoglobulin gene locus. These data shed light on a novel function of SETX in regulating the humoral immune response.

## Materials and Methods

### Ethics statement

Mouse experiments were approved by the McMaster University Animal Research Ethics Board. Human peripheral blood mononuclear cells (PBMCs) were obtained from healthy consenting donors, in accordance with a protocol approved by the Hamilton Integrated Research Ethics Board.

### Mice

SETX mice were a generous gift from Dr. Martin Lavin. SETX^+/+^ and SETX^-/-^ mice were genotyped using the methods described in Becherel *et al.* (30). Mouse experiments were performed using 6-12 week-old mice.

### Cells and Viruses

HeLa cells were originally obtained from the American Type Culture Collection (ATCC). Cells were maintained in Dulbecco’s Modified Eagle Medium (DMEM) supplemented with 2 mM GlutaMax (Gibco, Life Technologies), 10% fetal bovine serum (FBS) (Gibco, Life Technologies), 100 U/mL penicillin and 100 μg/mL streptomycin (Gibco, Life Technologies) at 37 °C with 5% CO_2_. Mouse embryonic fibroblasts (MEFs) were isolated from embryos at 13.5 days post-conception (35) and were maintained as described above, but with 15% FBS. Splenocytes, CH12 cells, and primary B cells were cultured in complete RPMI-1640 (cRPMI-1640), 2 mM glutamine, 10% FBS, 100 U/mL penicillin, 100 μg/mL streptomycin, and 10 mM β-mercaptoethanol (MilliporeSigma) at 37°C with 5% CO_2_. CH12-WT and CH12-SETX KO cells were a kind gift from Dr. Uttiya Basu (22). The Influenza A virus (IAV) strain used herein was A/Puerto Rico/8/1934 (IAV) which was propagated in 8-10 day-old embryonated chicken eggs (Canadian Food Inspection Agency) (29).

### Virus Inactivation

1 mg/mL of PR8 suspension was combined with 100× inactivation solution (0.925% formaldehyde in phosphate buffered saline (PBS)) to a final concentration of 1× inactivation solution. Following incubation at 4 °C on a rotator for 72 h, the inactivated virus was stored at −80 °C.

### Recombinant hemagglutinin (HA) production

To assess HA specific antibodies, recombinant PR8 HA was produced in HighFive cells as described in Margine *et al.*(36).

### Vaccination

Mice were immunized intramuscularly (*i.m.*, quadricep muscle) with 100 µL of 0.5 mg/mL formalin-inactivated PR8. Blood was drawn at 7, 14, and 28 days post-vaccination. Serum was separated by incubating blood overnight at 4 ℃ and centrifuged at 16,000 ×g for 5 min. Serum was frozen at −20 ℃ prior to further analyses.

### Infection

Mice were infected intranasally (*i.n.*) with 0.1 LD_50_ of PR8 (200 plaque forming units (PFU)). Mice were monitored for clinical signs and weight loss daily, with 80% of initial weight considered humane endpoint, in accordance with institutional guidelines. Blood was collected at 7- and 13- days post-infection. Serum was separated and frozen as mentioned above.

### Spleen and B cell isolation

Mice were euthanized via cervical dislocation and spleens were removed. Following removal, the spleen was homogenized using the Bullet Blender 24 Gold (Next Advance) and passed through a 100 μm filter. Cells were centrifuged at 200 ×g for 5 min at 4 °C and the supernatant was removed. RBCs were then lysed using 2 mL of ACK lysis buffer for 2 min at room temperature (RT). Following lysis, 40 mL of PBS was added to dilute the ACK and cells were centrifuged at 200 ×g for 5 min at 4 °C. The supernatant was removed, and the cells were resuspended in 1 mL of PBS containing 2% FBS and 1 mM EDTA. B cells were isolated from splenocytes using the EasySep Mouse B cell isolation kit (STEMCELL Technologies) according to the manufacturer’s instructions.

### B cell stimulation

For general polyclonal stimulation, cells were treated with 10 μg/mL lipopolysaccharide (LPS, MilliporeSigma), 1 μg/mL R848 (Invivogen), 1 μg/mL pokeweed mitogen (MilliporeSigma), *Staphylococcus aureus* Cowan (SAC, 1:10000) (MilliporeSigma), and 0.5 mM β-mercaptoethanol (MilliporeSigma) in cRPMI-1640. For IgA stimulation, cells were treated with 1 μg/mL CD40 (eBioscience), 20 ng/mL IL-4 (Peprotech), 1 ng/mL TGF-β (R&DSystems), and 0.5 mM β-mercaptoethanol (MilliporeSigma) in cRPMI-1640.

### Cell proliferation assay

B cells were isolated as described above and were stained with 10 μM carboxyfluorescein succinimidyl ester (CFSE) for 15 min at 37 °C. Following staining, cells were centrifuged at 200 ×g for 5 min at 4 °C and resuspended in cRPMI-1640. Cells were then plated in a 6 well plate and stimulated with 2 mL of polyclonal stimulation (described above) for 5 days. 72 hours after stimulation, the cells were collected and assayed with flow cytometry on the BD LSRFortessa (BD Biosciences) with at least one million events. B cells were quantified for the percentage of cells per cellular division using the proliferation tool in FlowJo v.10 (FlowJo, LLC, BD Biosciences).

### ELISpot

ELISpot assays were performed in 96 well MultiScreen Filter Plates (Millipore). Plates were coated with 50 μL of either capture antibody (goat anti-mouse IgG, IgA, IgM; ThermoFisher Scientific), or recombinant HA at 10 μg/mL for 24 h at 4 ^°^C in phosphate buffered saline (PBS). Plates were washed 3× with PBS with 0.1% Tween (PBS-T) and blocked with cRPMI-1640 for 1 h at 37 °C. Splenocytes were plated in a 96-well plate and serially diluted in cRPMI-1640 by 2-fold. The last column was left as a blank negative control. Plates were incubated overnight at 37 °C. Cells were removed and the plate was washed 3× with PBS-T. Biotinylated secondary antibodies, goat anti-mouse -IgM, -IgG, and -IgA (SouthernBiotech), at a 1:1000 dilution, were diluted in cRPMI-1640 and were incubated on plates for 1 h at RT. Secondary antibodies were aspirated and plates were washed 3× with PBS-T. Streptavidin Alkaline Phosphatase (SouthernBiotech), at a 1:3000 dilution, were diluted in cRPMI-1640 and were incubated on plates for 1 h at RT. Streptavidin Alkaline Phosphatase was aspirated and plates were washed 3× with PBS-T. BCIP/NBT substrate (ThermoFisher Scientific) was incubated on the plate for 20 min at RT. BCIP/NBT substrate was discarded and the plates were washed with distilled water before being left at RT overnight to dry. Plates were read using an ImmunoSpot plate reader (Image Acquisition 4.5) and the number of spots were analyzed using ImmunoSpot 3 (ImmunoSpot).

### Enzyme-linked immunosorbent assay (ELISA)

ELISAs were performed in 96-well NUNC-MaxiSorp™ plates (ThermoFisher Scientific). Plates were coated overnight at 4 ^°^C with 50 μL of either capture antibody (goat anti-mouse IgG, IgA, IgM; ThermoFisher Scientific), or recombinant HA at 5 μg/mL in bicarbonate/carbonate coating buffer (5.3 g Na_2_CO_3_, 4.2 g NaHCO_3_, pH 9.4, in 1 L of H_2_O). Plates were blocked using 100 μL of reagent and blocking diluent (0.5% BSA, 0.02% NaN_3_ in tris-tween wash buffer) for 1 h at RT. Serum samples were added and serially diluted 2-fold across the plate starting at a dilution of 1:640 for serum, leaving the last column as a blank control. Samples were then incubated for 1 h at 37°C. Following incubation, plates were washed 3× with TBS-T and 50 μL of goat anti-mouse-biotin - IgM (1:5000), -IgG (1:5000), and -IgA (1:2000) antibodies (SouthernBiotech), was added to reagent and blocking diluent, and incubated for 1 h at 37°C. Following incubation, plates were washed 3× with TBS-T and 50 μL of streptavidin-alkaline phosphatase (1:3000) was added to reagent and blocking diluent in each well and the plate was incubated for 1 h at 37°C. Following incubation, plates were washed 3× with TBS-T and 100 μL of pNPP one component microwell substrate solution was added for 10 min, following which the reaction was stopped with 100 μL of 3 N NaOH. Plates were then analyzed on the BioTek Synergy H1 plate reader (Agilent) at an absorbance of 405 nm. Endpoint titers were defined by the lowest dilution at which the O.D. was three standard deviations above the mean of the blank wells.

### Live/dead cell analysis

Isolated B cells received polyclonal stimulation as previously detailed for 3 or 5 days. At each time point, cells were stained with ReadyProbes cell viability imaging kit (ThermoFisher Scientific) according to the manufacturer’s protocol. Following staining, cells were imaged using the EVOS-FL system (ThermoFisher Scientific) at 10× magnification. The percentage of dead cells was quantified by overlaying the live (blue) and dead (green) channels and counting the proportion of cells which exhibited dual staining.

### Immunofluorescence

MEFs and HeLa cells were cultured overnight on coverslips and treated with either 0.5 nM H_2_O_2_ (MilliporeSigma) or 20 μM phleomycin (Invivogen). At the indicated time points, cells were fixed using ice-cold methanol for 20 min at −20 °C. Following methanol fixation, cells were washed 3× with PBS. Cells were then permeabilized using 0.2 % Triton X/ 0.1 % BSA diluted in PBS for 15 min at RT, and then washed 3× PBS. Following permeabilization, cells were blocked with 10 % normal goat serum (Abcam)/1 % BSA diluted in PBS for 30 min. Cells were incubated with primary antibodies overnight at 4 °C. Primary antibodies included rabbit anti-human SETX (1:100; Bethyl Laboratories, Inc.), mouse anti-mouse/human γH2AX (1:1000; Abcam), mouse anti-human 53BP1 (1:100; MilliporeSigma), mouse anti-mouse/human Phospho-ATM (1:200; ThermoFisher Scientific), and mouse anti-R-loop S9.6 (1:100; Kerafast). Cells were then washed 5× with PBS and probed with compatible AlexaFluor-conjugated secondary antibodies at a 1:1000 dilution (Life Technologies). Cells were washed 3× with PBS and DNA was counterstained using Hoechst 33342 (1 μg/mL) (Life Technologies) and mounted with EverBrite mounting medium (Biotium). Images were taken with the Zeiss Imager M2 microscope (Carl Zeiss Canada Ltd.) at the indicated magnification. For DNA damage kinetics experiments, cells were considered positive for p-ATM and ɣ-H2AX foci formation if the number of foci was greater than the average number of p-ATM/ɣ-H2AX foci in the SETX^+/+^ untreated conditions + 3 standard deviations.

### Immunoprecipitation (IP)

Human PBMCs were isolated from peripheral blood and collected into ethylenediamine tetra-acetic acid (EDTA) coated tubes (BD Biosciences). PBMCs were processed using density gradient centrifugation. In short, 6 mL of blood, 3 mL of RT Histopaque 1119 (MilliporeSigma), and 3 mL of RT Histopaque 1077 (MilliporeSigma) were successively layered on top of each other and centrifuged at 930 ×g for 30 min at 4 °C with no deceleration. The PBMC layer was isolated and washed in PMN buffer (0.5% BSA, 0.3 mM EDTA in HBSS) twice and centrifuged at 450 ×g for 5 min at 4 °C. PBMC’s were stimulated with polyclonal stimulation, as described above, for 72 h at which point crude lysates were collected using NP-40 lysis buffer (0.15 M NaCl, 0.05 M TRIS (pH 8.0), and 1% NP-40) with Halt Protease and Phosphatase Inhibitor Cocktail (100X) (ThermoFisher Scientific). Protein amounts were quantified using the Pierce BCA protein assay kit (ThermoFisher Scientific) according to the manufacturer’s instruction. Prior to IP, 500 μg of total protein was precleared with protein G agarose beads (Invitrogen) overnight at 4 °C with constant agitation. Following preclearing, samples were centrifuged at 16,000 ×g for 5 min at 4 °C and the protein was incubated with 10 μg of either SETX (Bethyl Laboratories Inc.) or control IgG (Abcam) antibody and incubated overnight at 4 °C with constant agitation. Samples were centrifuged at 16,000 ×g for 5 min at 4 °C and washed 10× with lysis buffer, with samples being centrifuged at 16,000 ×g for 5 min at 4 °C between washes. Protein was eluted from bead/protein complexes using 2× Pierce LDS sample buffer (ThermoFisher Scientific) with 5% β-mercaptoethanol at 95 °C for 10 min.

### Western blotting

Protein was loaded onto a 4-12% Bis-Tris gel (ThermoFisher Scientific) and ran at 100V for 90 min with Bolt MOPS SDS running buffer (Invitrogen). Afterwards, protein was transferred onto a PVDF membrane using 1x Towbin transfer buffer (0.025 M TRIS, 0.192 M glycine, pH 8.6), containing 3% methanol, for 18.5 h at 4 °C. The membrane was blocked in 5% skim milk in Tris-buffered saline with 0.05% Tween (TBS-T) at RT for 4 h. The membrane was then incubated with rabbit anti-SETX (1:1000; Bethyl Laboratories, Inc.), rabbit anti-53BP1 (1:1000; Abcam), or mouse anti-GAPDH (1:3000; ProteinTech) antibodies, diluted in 5% milk in TBS-T overnight at 4 °C. Membranes were then washed 5× with TBS-T and subsequently incubated with anti-mouse or anti-rabbit IgG-HRP conjugated antibodies (ThermoFisher Scientific) diluted 1:4000 in 5% skim milk in TBS-T for 1 h at RT. Following incubation, the membranes were washed 5× with TBS-T and then were given a final wash with TBS. Membranes were imaged using Pierce ECL Western Blotting Substrate (32106, Invitrogen) and filmBlu-Lite Autoradiography film (DIAFILM810-LITE, Diamed).

### Germline transcript quantification

RNA was isolated from stimulated mouse splenocytes using the RNeasy isolation kit (QIAgen) and 1 μg of complimentary DNA was synthesized using the Maxima First Strand cDNA Synthesis kit (ThermoFisher Scientific) according to the manufacturer’s instructions. Expression levels of AID, IgM GLT (M-GLT), and IgA GLT (A-GLT) were assessed using the SensiFAST SYBR reaction kit (Bioline) and quantitative PCR (q-PCR) along with previously published primers (37). The cycle threshold (Ct) was obtained for each probe and normalized to a house keeping gene (ΔCt) followed by normalization to the value of the first probe (ΔΔCt).

### DNA:RNA Immunoprecipitation (DRIP)

DRIP was performed using the methods described in Halász *et al.* as a framework(38). Briefly, DNA was isolated from stimulated mouse splenocytes using the DNeasy isolation kit (QIAgen), according to the manufacturer’s protocol. The total amount of DNA was quantified, and DNA was sonicated to a size ranging from 250-1000 bp. 6 μg of total DNA was then precleared using protein G agarose beads (Invitrogen) for 1 h at 4 °C while shaking, following which samples were either treated with RNaseH (New England Biolabs) or left untreated. Samples were incubated with 10 μg of S9.6 antibody (Kerafast), or 10 μg of control mouse IgG (Invitrogen), while shaking overnight at 4 °C. Protein G agarose beads (Invitrogen) were then added to the samples and incubated at 4 °C while shaking overnight. The protein G agarose beads were then spun at 16,000 ×g for 5 min and washed 4× with buffers of differing ionic strength. The first wash, low salt IP buffer (16.7 mM TRIS (pH 8), 1.2 mM EDTA, 167 mM NaCl, 0.01% SDS, and 1.1% Triton X-100), the second wash, high salt IP buffer (16.7 mM TRIS (pH 8), 1.2 mM EDTA, 500 mM NaCl, 0.01% SDS, and 1.1% Triton X-100), the third wash, high LiCl buffer (10 mM TRIS (pH 8), 1 mM EDTA, 250 mM LiCl, and 1% NP-40), and the final wash, low LiCl buffer (10 mM TRIS (pH 8), 1 mM EDTA, and 50 mM LiCl). Following the washes, DNA was eluted from protein G agarose beads (Invitrogen) using 100 μL elution buffer (50 mM TRIS (pH 8), 10 mM EDTA, and 1% SDS) at 65 °C for 15 min. After the 15 min incubation, an additional 100 μL of PBS was added, and the DNA was purified using the DNeasy isolation kit (QIAgen). Following DNA isolation, samples were assessed via quantitative PCR (qPCR) using SensiFAST SYBR reaction kit (Bioline) and previously published primers against areas of the IgH locus(37).

### High throughput immunoglobulin repertoire sequencing

High throughput-sequencing analysis of mouse immunoglobulin repertoire was performed as previously described(39). Briefly, SETX^+/+^ and SETX^-/-^ mice were vaccinated *i.m.* with formalin inactivated-PR8 and spleens were isolated 14 days post-vaccination. Total RNA was isolated using the RNeasy Qiagen RNA isolation kit, according to the manufacturer’s protocol. 1 μg of total RNA was used for the sequencing of the IgH repertoire, according to the Turchaninova *et al.* protocol(39). Illumina adaptors were added to the library according to the manufacturer’s protocol.

Sequencing was performed on an Illumina Mi-Seq platform using asymmetric 400+200-nt paired-end sequencing. The MIGEC software was used to demultiplex the samples and perform UMI-based assembly of full length V regions (40). ChangeO package was used to perform IgBLAST and perform clonal clustering (41). Immunoglobulin gene usage analysis was conducted using the Alakazam package (41, 42). The ShazaM package was used to quantify mutations within the full V-region or in the CDR/FDR regions (41, 43).

### Microhomology analysis

DNA was isolated from SETX^+/+^ and SETX^-/-^ CH12 cells either treated with IgA-specific stimulation or non-stimulated controls. PCR was run using unique barcoded primer sets for each condition. DNA samples were then purified using AMPure Beads PB (Pacific Biosciences), pooled in equimolar concentrations, purified again with AMPure Beads PB, and sent to the Mobix Lab (McMaster University) for Single Molecule, Real-Time (SMRT) sequencing (Pacific Biosciences).

PacBio reads were aligned to the mouse genome GRCm38 using the minimap2 (v2.21)(44). Duplicate reads were marked using and removed using SAMtools (v1.7)(45). Structural variations were detected using sniffles (v1.0.12) (46). Only deletions that at least partially covers the switch region (chr12: 113224585 – 113389342), have more than 10 supporting reads, and with the quality filter “PASS” were considered. Repair pathways for each deletion were categorized with the following criteria as previously described (47): number of homology base pairs < 2 were considered “NHEJ”, 2 ≤ number of homology base pairs ≤ 20 were considered “MMEJ”, 20 < number of homology base pairs ≤ 50 were considered “SSA”, and number of homology base pairs > 50 were considered “HR”.

### Statistical Analysis

All data is represented using GraphPad Prism (v10.4.1) (GraphPad Software) and statistical analysis was performed using GraphPad Prism.

## Results

### Senataxin interacts with DDR proteins involved in CSR

The DDR is known to play a prominent role in CSR including repairing the DSBs that are formed during CSR to ensure proper switching of immunoglobulin isotypes (5–7). Indeed, SETX has been previously demonstrated to colocalize with multiple markers of the DDR in HeLa cells, many of which have essential functions during CSR (32). Therefore, we hypothesized that SETX might play a role in the DDR induced in B cells during CSR. We first confirmed that SETX colocalizes with Tumor Suppressor p53-Binding Protein 1 (TP53BP1 or 53PB1) and Ser^139^ Phosphorylated H2A Histone Family Member X (γ-H2AX) in HeLa cells using immunocytochemistry (Supplemental Fig. 1A). Subsequently, we demonstrated that SETX colocalizes with R-loops in HeLa cells both under physiological conditions and following DNA damage induced by phleomycin – an intercalating agent that induces double-strand breaks (Supplemental Fig. 1A). To explore this interaction in a more physiologically relevant model system, we used human primary B cells to confirm the interaction of SETX with DDR proteins by immunoprecipitation. We observed that 53BP1 coimmunoprecipitates with SETX upon stimulation of CSR (Supplemental Fig. 1B).

### Senataxin deficiency alters the kinetics of the DDR

Having confirmed that SETX interacts with DNA damage repair proteins, we explored the role of SETX on the kinetics of the DDR. Since this is difficult to monitor using immunofluorescence techniques in B cells due to their morphology non-adherent nature, we utilized mouse embryo fibroblasts (MEFs) treated with various DNA damaging agents. SETX^+/+^ and SETX^-/-^ MEFs were treated for 30 min with 0.5 mM H_2_O_2_ – which acts as a general DNA damaging agent – or 20 μM phleomycin – which induces double-strand breaks. We tracked γ-H2AX foci (Figs. 1A and 1B) and phosphorylated ataxia-telangiectasia mutated (p-ATM) foci (Figs. 1C and 1D) using immunofluorescence before the addition of the DNA damaging agent as well as at 0 h, 0.5 h, 1 h, 2 h, 4 h, 8 h, and 24 h after removal of the DNA damaging agent.

**Figure 1:**
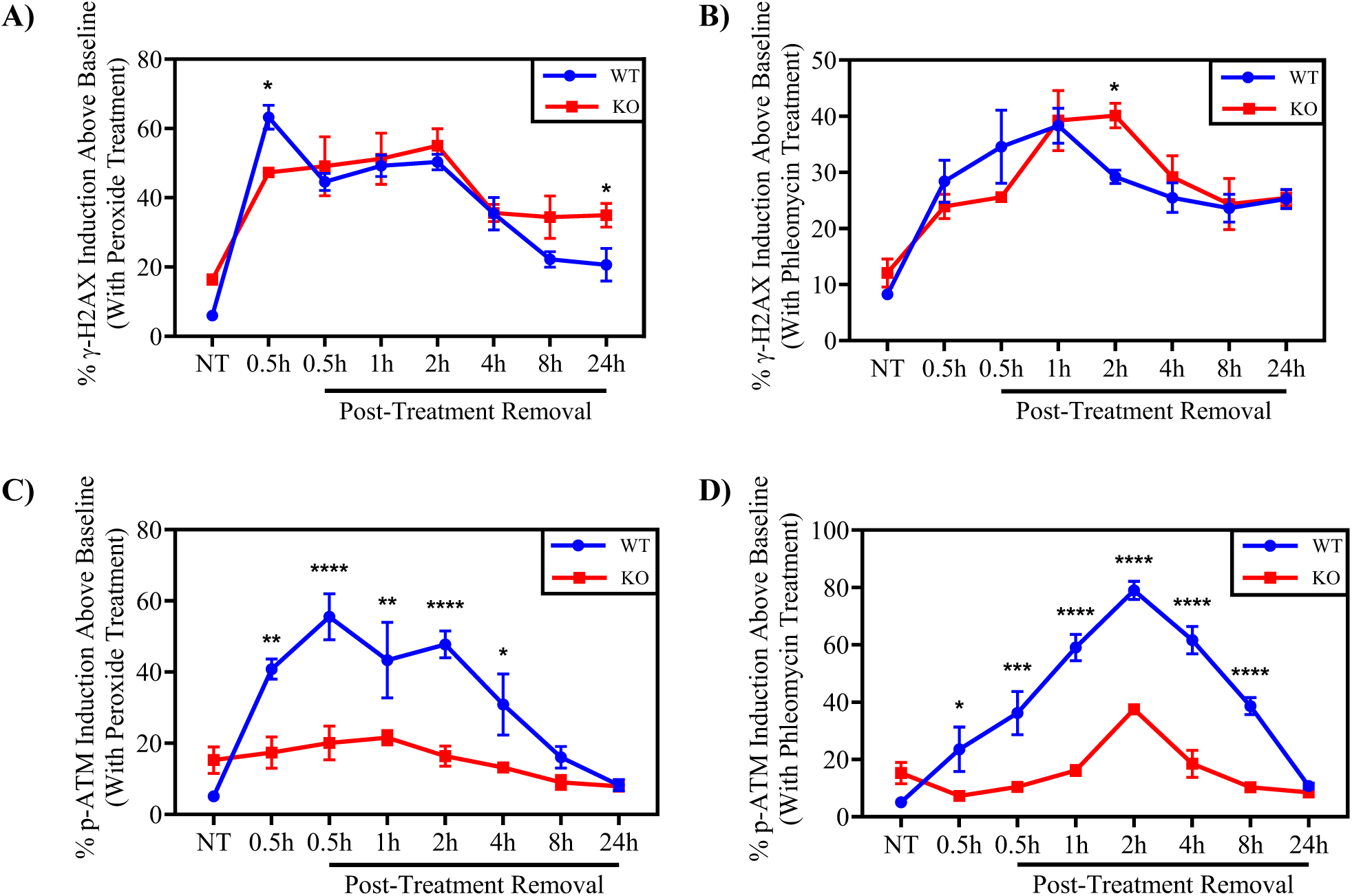
SETX alters the induction of p-ATM foci but not γ-H2AX foci following DNA damage. SETX^+/+^ (WT) and SETX^-/-^ (KO) MEFs were treated with 0.5 mM H_2_O_2_ or 20 µM phleomycin for 30 min, following which the damaging agent was removed and replaced with fresh media and cells were fixed with 3.7% PFA at 30 min, 1 hr, 2 hrs, 4 hrs, 8 hrs, and 24 hrs post removal of the damaging agent. Immunofluorescence for the common DNA damage markers γ-H2AX and p-ATM was performed at each time point. **(A and B)** In order to quantify the level of DNA damage in the MEFs, the number of γ-H2AX foci/cell in the untreated SETX^+/+^ MEFs were averaged and the average number of foci plus three standard deviations was considered as positive for induction of γ-H2AX and compared SETX^+/+^ MEFs to SETX^-/-^ MEFs treated with either **(A)** H_2_O_2_ or **(B)** phleomycin. **(C and D)** Quantification of p-ATM induction levels following treatment with **(C)** H_2_O_2_ or **(D)** phleomycin over the indicated time course. Statistics were performed on n = 3 with p < 0.05, *, p < 0.01, **, p < 0.001, ***, and p < 0.0001, **** determined via two-way ANOVA with Tukey post-hoc test.

Following treatment with H_2_O_2_ we observed significant differences in γ-H2AX induction when comparing SETX^+/+^ and SETX^-/-^ MEFs at 0.5 h post-treatment and 24 h post-treatment removal (Fig. 1A). Upon treatment with phleomycin, we observed significant differences in γ-H2AX induction between SETX^+/+^ and SETX^-/-^ MEFs at 2 h post-treatment removal (Fig. 1B). H2AX foci were induced in both SETX^+/+^ and SETX^-/-^ MEFs when compared to baseline (Figs. 1A and 1B). In addition to γ-H2AX, we assessed the effect of SETX deficiency on p-ATM foci formation after inducing DNA damage as ATM responds to DSB induction and is rapidly auto-phosphorylated upon activation (48). Upon treatment with H_2_O_2_, SETX^+/+^ MEFs responded strongly and rapidly to DNA damage, inducing high levels of p-ATM foci positive cells whereas SETX^-/-^ MEFs exhibit minimal induction of p-ATM foci throughout the course of the experiment (Fig. 1C).

Likewise, upon treatment with phleomycin, p-ATM foci formation was potently induced in SETX^+/+^ MEFs, while p-ATM foci induction was much lower in the SETX^-/-^ MEFs (Fig. 1D). Again, both SETX^+/+^ and SETX^-/-^ MEFs significantly induce p-ATM when compared to baseline (Figs. 1C and 1D). Together, these results suggest that SETX^-/-^ cells display an abnormal DDR, and in particular, have a significant defect in recruiting factors that are involved in recognizing and/or repairing DNA damage.

### SETX deficiency alters the magnitude and rate of class switch recombination ex vivo

Given the role of both the DDR and the resolution of R-loops in CSR (5–7), we sought to explore the role of SETX on CSR. To this end, splenocytes were isolated from spleens of naïve SETX^+/+^ and SETX^-/-^ mice and stimulated *ex vivo* with polyclonal stimulants to induce CSR (49). Following stimulation, ELISpots were used to assess the frequency of IgM-, IgG- and IgA-secreting B cells at days 0, 1, 3, and 5 post-stimulation (Figs. 2A, 2B, and 2C). Frequencies of both IgM- and IgG-producing B cells were similar throughout the time-course of stimulation between SETX^+/+^ and SETX^-/-^ mice (Figs. 2A and 2B). However, 5 days post-stimulation, we observed a significant decrease in the number of IgA-producing B cells from SETX^-/-^ mice relative to SETX^+/+^ mice (Fig. 2C). Indeed, the frequency of IgA-producing B cells from SETX^-/-^ mice remained relatively unchanged over the 5 days post-stimulation, while the frequency of IgA-producing B cells from SETX^+/+^ mice increased (Fig. 2C). These results demonstrate that SETX deficiency in B cells display causes impaired IgA class-switching upon *ex vivo* polyclonal/antigen-independent stimulation.

**Figure 2:**
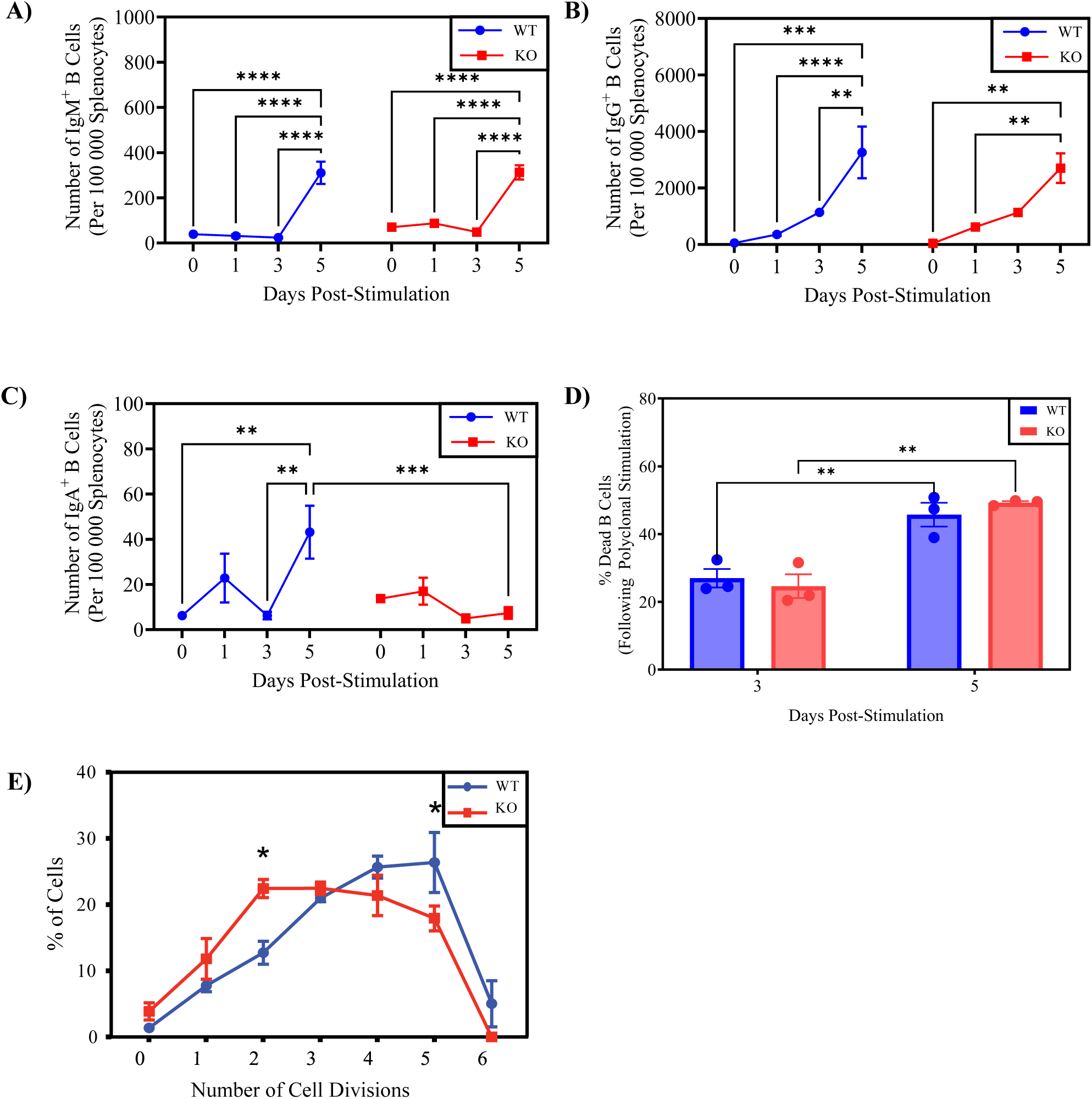
SETX deficiency alters the magnitude and kinetics of class-switch recombination *ex-vivo* without altering cell death in B cells. SETX^+/+^ (WT) and SETX^-/-^ (KO) naïve splenocytes were isolated and stimulated with a polyclonal stimulation. Antibody isotype switching was observed following 0, 1, 3, and 5 days post-stimulation and demonstrated as **(A)** IgM, **(B)** IgG, and **(C)** IgA isotypes. n = 3 **(D)** B cells were stained with live/dead stain at days 3 and 5 post-stimulation. The percentage of dead cells were quantified by the overlap of the live and dead staining of cells. n = 3. **(E)** B cells were stained with 10 μM CFSE prior to stimulation and stimulated for 72 hrs. CFSE stained cells were assessed using flow cytometry with a minimum cell count of 1,000,000 cells. B cells were quantified for the percentage of cells per cellular division using the proliferation tool in FlowJo. n = 3. Statistical analysis was performed using two-way ANOVA with a Bonferroni post-hoc test. p< 0.05, *, p < 0.01, **, p < 0.001, ***, p < 0.001, ****

### SETX deficiency impacts B cell proliferation

Given that SETX deficiency impaired IgA class-switching *ex vivo,* we next explored whether these differences are due to differences B cell death or proliferation. Using a live/dead stain, we did not observe any significant differences in the percentage of dead SETX^+/+^ and SETX^-/-^ B cells at days 3 and 5 post-stimulation (Fig. 2D). Given that switching to IgA requires 5-7 rounds of cellular division (50), we also explored the rate of proliferation between SETX^+/+^ and SETX^-/-^ B cells as a possible explanation of the difference in IgA-secreting B cell frequencies. We chose 72 h post-stimulation for our assessment as SETX^+/+^ and SETX^-/-^ B cells diverge in IgA expression at 5 days post-stimulation and thus, proliferation deficits were likely to manifest at earlier times. 72 h post-stimulation polyclonal stimulation, SETX^-/-^ B cells had a higher proportion of cells having completed 0-2 cell divisions as well as a decrease in the percentage of cells having undergone a great number of divisions (4–6) when compared to SETX^+/+^ B cells (Fig. 2E). Consequently, we observed that a SETX deficiency results in proliferative defects in B cells post-stimulation but not increased cell death.

### SETX deficiency does not affect IgA production in response to vaccination

Given the deficiency in IgA-secreting SETX^-/-^ B cells observed *ex vivo,* we subsequently explored the profile of serum antibodies following vaccination to determine if this deficiency was still found *in vivo* in the presence of the additional cell types required for CSR. To assess the role of SETX in a natural model of antigen-dependent CSR, SETX^+/+^ and SETX^-/-^ mice were vaccinated intramuscularly (*i.m.*) with formalin-inactivated PR8 H1N1 influenza A virus (IAV) and the serum titers of both total and antigen-specific IgM, IgG, and IgA antibodies were analyzed at days 7, 14, and 28 post-infection (Figs. 3A-C). At all time points, the total and antigen-specific antibody titers for IgA, IgG, and IgM remained equivalent when comparing SETX^+/+^ and SETX^-/-^ mice (Figs. 3A-C). As such, these results demonstrate that SETX^-/-^ mice do not exhibit a deficiency in antibody titers during a primary immune response to vaccination.

**Figure 3:**
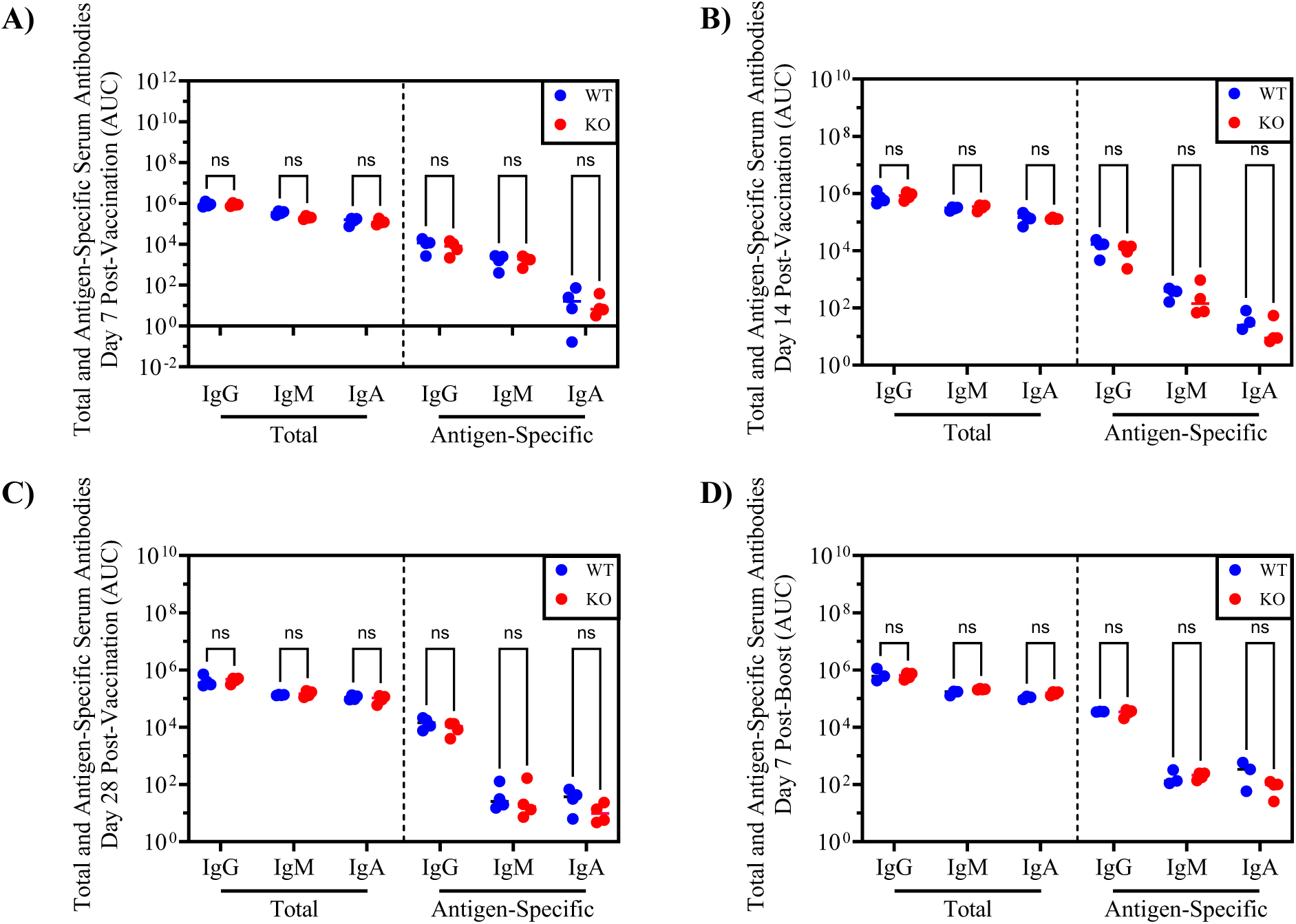
SETX deficiency does not alter the primary or secondary immune response to vaccination. SETX^+/+^ and SETX^-/-^ mice were vaccinated with 100 µL of 0.5 mg/mL formalin-inactivated PR8 influenza A virus *i.m..* Following vaccination, we performed cheek bleeds on days 7, 14, and 28. Serum was isolated from the blood samples and via ELISA we quantified both total and PR8-HA specific antibody isotype levels. The endpoint titers were plotted for **(A)** day 7, **(B)** day 14, and **(C)** day 28 post-vaccination (n = 3 – 5). **(D)** A secondary immune response was elicited via homologous boost at 28 days post-vaccination and antibody levels in sera were assessed via ELISA 7 days post-boost (n = 3 – 4). Statistical analysis was performed using multiple t-tests with correction for multiple comparisons using the Holm-Šídák method.

To explore whether a SETX deficiency affected IgA production during a memory response, we performed a homologous boost 28 days after the first infection. At day 7 post-boost, there was no significant difference in the total or antigen-specific antibody titers of IgA, IgM, or IgG antibodies between the SETX^+/+^ and SETX^-/-^ mice (Fig. 3D). These results suggest that a SETX deficiency does not affect the reactivation of previously switched IgA B cells.

### SETX deficiency impacts IgA antibody titers mounted in response to infection

We went on to study how SETX regulates the antibody response to natural infection. SETX^+/+^ and SETX^-/-^ mice were infected intranasally (*i.n.*) with 0.1 LD_50_ (250 PFU) of PR8 H1N1 influenza A virus (IAV). Serum titers of both total and antigen-specific IgM, IgG, and IgA antibodies were analyzed at days 7 and 13 post-infection (Figs. 4A, B). We did not observe any major differences in total antibody titers at day 7 or 13 post-infection in serum (Figs. 4A, B). While no differences in antigen-specific antibody titers were observed at day 7 post-infection in serum (Fig. 4A), significantly lower antigen-specific IgA titers – but not IgG or IgM – were found in SETX deficient mice when compared to WT mice (Fig. 4B). Thus, SETX^-/-^ mice mount reduced antigen-specific IgA titers in the response to viral infection.

**Figure 4:**
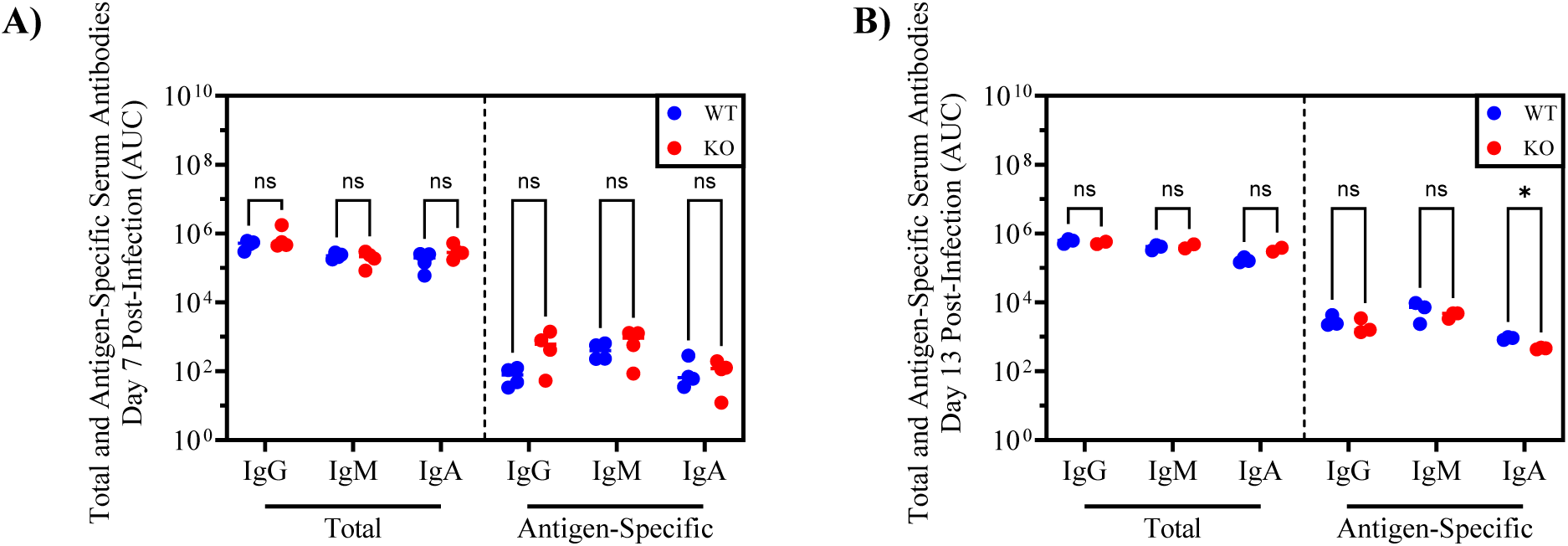
SETX deficiency alters the primary immune response to viral infection. SETX^+/+^ and SETX^-/-^ mice were infected with 200 PFU of PR8 influenza virus *i.n.*. Following infection, we performed cardiac bleeds on days 7 and 13. Serum was isolated from the blood samples. Using ELISA, we quantified both total and PR8-HA specific antibody isotype levels in serum. The endpoint titers were plotted for serum at **(A)** day 7 (n = 4) and **(B)** day 13 (n = 3) post-infection. Statistical analysis was performed using multiple t-tests with correction for multiple comparisons using the Holm-Šídák method., p < 0.05, *.

### SETX deficiency reduces the diversity of the IgA repertoire post-vaccination

Although we observed no difference in antigen-specific IgA titers following either vaccination or viral infection *in vivo*, we next specifically characterized IgA repertoire diversity in SETX deficient conditions. To explore diversity within the IgA repertoire, SETX^+/+^ and SETX^-/-^ mice were vaccinated *i.m.* with formalin-inactivated IAV and spleens were isolated 14 days post-vaccination. To assess the diversity of the IgH repertoire we used a molecular barcoding, high quality sequencing approach (39). The IgH repertoire was visualized using Circo plots for both SETX^+/+^ (Fig. 5A) and SETX^-/-^ (Fig. 5B) mice. All sequences on the bottom half of the circle were mapped to their respective antibody isotype, on the top half of the circles. Additionally, the size of the connecting, coloured lines represent the number of reads that mapped to that sequence.

**Figure 5:**
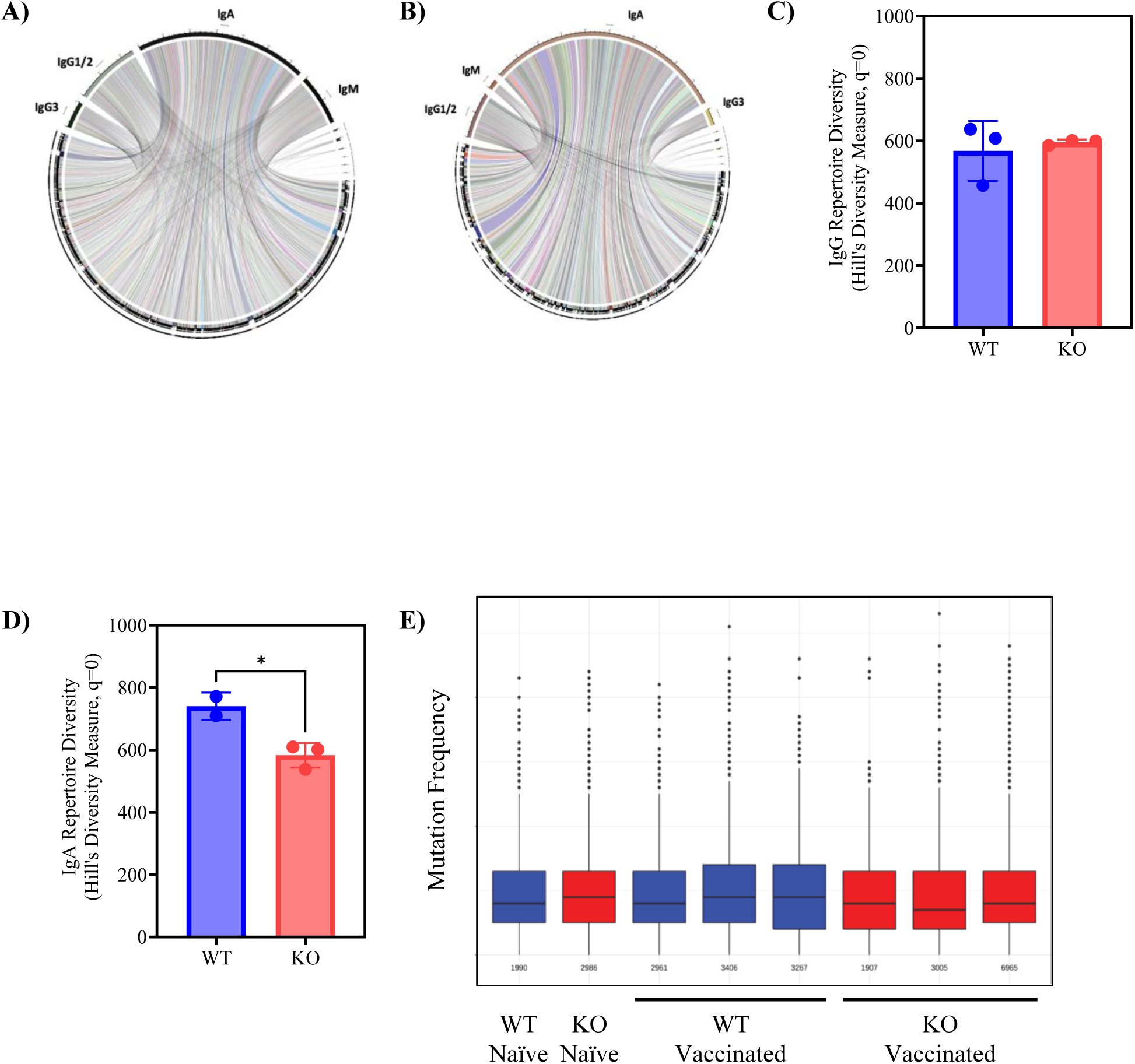
SETX deficiency alters the diversity within the IgA repertoire. SETX^+/+^ (WT) and SETX^-/-^ (KO) mice were vaccinated with formalin-inactivated IAV *i.m.*. 14 days post-vaccination, spleens were removed, and RNA isolated from whole splenocytes**. (A and B)** The Ig repertoire was assessed via deep-sequencing and diversity was visualized using circo plots, **(A)** WT and **(B)** KO. **(C and D)** The Hills diversity of the **(C)** IgG and **(D)** IgA repertoire was quantified at q = 0. Statistics were performed on n = 3 with p < 0.05, * determined via Student’s t-test. **(E)** The WT and KO mice IgA variable regions were assessed for the overall mutation rate compared to a germline sequence (n = 3).

Quantification of both the IgA and IgG repertoire diversity was calculated using the Hill’s diversity measure with q = 0. The IgG repertoire demonstrated no difference in diversity between SETX^+/+^ and SETX^-/-^ mice (Fig. 5C), whereas the SETX^-/-^ IgA repertoire was significantly less diverse than the SETX^+/+^ IgA repertoire (Fig. 5D). Interestingly, mutation rates within the IgA repertoire (Fig. 5E) were unaffected, suggesting that a SETX deficiency does not affect somatic hypermutation (SHM) rates. These results suggest that fewer B cells are filling the IgA repertoire in SETX deficient conditions, indicating a difficulty in class switching to the IgA isotype.

### SETX deficiency alters the distribution of R-loops within the IgH locus

As SETX acts directly upon R-loops as an RNA:DNA helicase (25, 28), we aimed to quantify the relative abundance of R-loops at specific regions of the IgH locus (Fig. 6A). To this end, B cells received either polyclonal stimulation, IgA-specific stimulation, or no treatment for 24 h. DNA isolation and R-loop immunoprecipitation (DRIP) was then performed on the treated and control B cells. Samples were treated with RNase H as a negative control to remove R-loops and no signal was observed following RNase H treatment (data not shown), as expected. We observed that a SETX deficiency results in an increase in the proportion of R-loops at different regions within the IgH locus (Figs. 6B-D). Polyclonal stimulation results in an increase in R-loops within both the Sμ and the Sα regions in SETX^-/-^ cells (Fig. 6C) with further elevation of R-loops levels upon IgA stimulation in SETX^-/-^ cells (Fig. 6D). Additionally, we quantified the relative expression levels of GLTs and found that the SETX^-/-^ cells have a trend towards increased GLT expression levels compared to SETX^+/+^ cells (Fig. 6E). These results suggest that the effect of the SETX deficiency on IgA class switching is downstream of transcription of the IgA locus.

**Figure 6:**
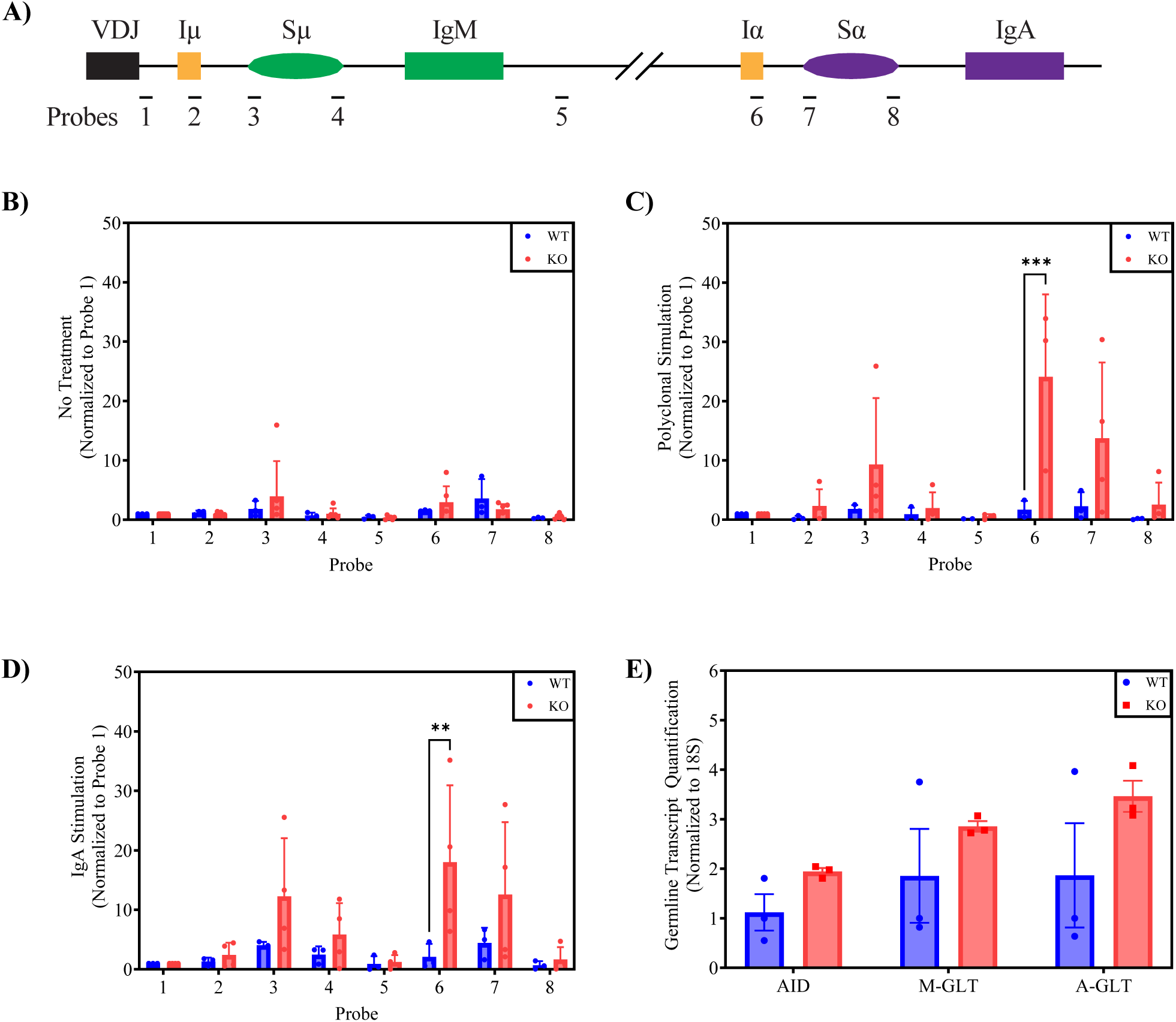
Senataxin deficiency alters the localization of R-loops within the IgH locus. To assess the role of SETX on R-loop formation within the IgH locus, we isolated R-loops from whole DNA and probed the IgH locus for their prevalence. **(A)** A not-to-scale representative image of the mouse IgM and IgA regions within the IgH locus, approximate probe locations are marked. DNA was isolated from whole splenocytes and R-loops were immunoprecipitated following 24 hours of **(B)** no treatment, **(C)** polyclonal stimulation, and **(D)** IgA specific stimulation. **(E)** RNA was isolated from splenocytes treated with IgA stimulation for 72 hours and the expression levels of AID, IgM-GLT, and IgA-GLT were assessed via q-PCR (n = 3). Statistics were determined via two-way ANOVA with Bonferroni post-hoc test, p < 0.001, ***.

### Senataxin does not alter the mechanism of the DDR during CSR

During CSR, DSBs are ultimately repaired using classical NHEJ(51). However, a secondary pathway, termed alternative end joining (a-EJ), can also repair DSBs when NHEJ is compromised. A key distinction between the two processes is that a-EJ takes longer than NHEJ, and a-EJ relies upon microhomology within DNA sequences to initiate repair of the DSBs (52–54). Our earlier findings indicate a delay in the kinetics of DNA repair and suggested that a-EJ may be more prevalent in SETX-deficient conditions. To determine the DNA repair pathway utilized, we used a CRISPR SETX knockout CH12 B cell line and sequenced recombined S-regions from IgA+ cells to analyze microhomology (Supplemental Fig. 2A). We isolated DNA from stimulated and unstimulated SETX^+/+^ and SETX^-/-^ CH12 cells and performed single molecule real time (SMRT) sequencing on the S-regions to analyze microhomologies. We observed no significant differences between the SETX^+/+^ and SETX^-/-^ CH12 cells in the utilization of alternative methods of DNA repair (Supplemental Fig. 2B) suggesting that there is no alteration in repair pathway choice in SETX^-/-^ B cells.

## Discussion

SETX has been associated with the DDR through interactions with the DDR markers 53BP1 and γ-H2AX in HeLa cells (30–33). Additionally, phleomycin treatment has previously been demonstrated to increase the number of SETX foci in HeLa cells through elevated DSB induction (32). The DDR and 53BP1 are integral for efficient CSR (55) and recently, 53BP1 has been shown to be important for the 3D reconstruction of the IgH locus during CSR (56, 57). Given that SETX colocalizes with DDR proteins following DNA damage, SETX may act as a scaffold linking 53BP1 to the IgH locus via SETX/R-loop interactions.

While SETX has been demonstrated to interact with DDR proteins, the effect of SETX on the kinetics of the DDR has not been explored. We observed that γ-H2AX foci induction was similar between SETX^-/-^ and SETX^+/+^ MEFs. In addition to γ-H2AX, we assessed the role of SETX on p-ATM activation using an antibody specific for ATM phosphorylation at site serine 1981; this site stabilizes ATM at the sites of DSB, allowing for the DDR and DNA repair to occur (48). Additionally, p-ATM phosphorylates γ-H2AX and anchors many of the proteins involved in the DDR (58, 59), including those known to interact with SETX during DNA damage. In contrast to γ-H2AX, p-ATM foci formation was dramatically delayed in SETX^-/-^ MEFs following DNA damage. p-ATM, ATR – another mediator of the DDR, and DNA-PK all phosphorylate H2AX (60, 61). Thus, in p-ATM-deficient states, as seen in SETX^-/-^ cells, other kinases allow for redundancy between activators of H2AX and the unaltered formation of γ-H2AX. The functional roles of ATM and γ-H2AX on the DDR suggests an issue with the appropriate activation of essential DNA repair factors through p-ATM. Specifically, while phosphorylation of H2AX is crucial for the recruitment of DDR factors (62), p-ATM is key regulator of the activity of DDR factors (58). Therefore, in SETX^-/-^ conditions, DDR factors may be recruited and but have altered functions resulting in abnormal DNA repair.

SETX is involved in many cellular functions that are essential for CSR, including R-loop resolution (23, 63), cytokine production (29), DNA double strand breaks repair (30), and the DDR (30, 32–34). RNA helicases have been demonstrated to be key contributors in CSR (22, 37), however, the role of SETX has only been tangentially explored. Although the infertility of male SETX^-/-^ mice is attributed to R-loop accumulation in the testes during meiosis and subsequent cell death (30), we observed no similar increase in cell death in SETX^-/-^ B cells. Interestingly, the delay in cell division seen in the SETX^-/-^ B cells is consistent with the lack of IgA observed following *ex vivo* stimulation of SETX^-/-^ cells as IgA switching requires several rounds of cell division (50).

Our *in vivo* vaccination of SETX^+/+^ and SETX^-/-^ mice did not result in a difference in the antibody response after primary or secondary exposure to antigen. However, we observed a significant decrease in IgA antibody titers 13 days after IAV infection. The use of a larger sample size and more sensitive assays, such as ELISpots (64), for exploring CSR after vaccination or infection may be needed to detect additional differences in CSR between SETX^+/+^ and SETX^-/-^ mice. Furthermore, previous studies have observed a difference in CSR between T-cell dependent and T-cell independent antigens including a preference for CSR to IgG with T-cell dependent antigen stimulation (65). Our vaccination and infection protocols stimulate a T-cell dependent response (66) compared to the T-cell independent response from *ex vivo* polyclonal stimulation (67). In conjunction with the lack of T-cells in our *ex vivo* stimulation, this may explain some of the difference between the *ex vivo* and *in vivo* data.

However, we did observe decreased diversity of the IgA repertoire in SETX deficient mice. Clinically, a selective deficiency in IgA, SIgAD, is the most common form of primary immunodeficiency (68). While typically asymptomatic, SIgAD has been associated with a potential increase in susceptibility to mucosal infections (68, 69) and a higher incidence of autoimmune disorders (70, 71). Although SETX^-/-^ mice do not exhibit SIgAD – as their total levels of IgA are comparable to those of SETX^+/+^ mice, the observed lack IgA repertoire diversity suggests a potentially unique phenotype of SIgAD in the context of AOA2.

Antigen specificity of B cells is gradually improved through a process known as affinity maturation and occurs in GCs. The amount of somatic hypermutation in antibody V-regions that drives affinity maturation typically increases with the number of cell divisions (72). We observed no difference in the magnitude of SHM within the variable regions of IgA-switched SETX^+/+^ and SETX^-/-^ B cells. Thus, while SETX deficiency alters CSR, we did not detect a role for SETX in SHM. Even though SHM and CSR share many similarities with regards to AID targeting and activity (73), CSR induces DSBs while SHM introduces mutations into the variable region independent of DSBs (14). Although the mechanistic differences between SHM and CSR are incompletely understood, our data suggests that SETX acts on CSR downstream of AID activity, potentially on the DSB DDR, as this is a point at which CSR and SHM differ.

As one of the major cellular roles of SETX is the resolution of R-loops (23, 25), SETX deficient cells exhibit elevated R-loops levels. R-loops are essential for the completion of CSR, especially within the S regions (74–76), and genetically altering B cells to prevent R-loop formation also inhibited CSR (74, 75). We found that R-loops were elevated across both the IgM and IgA regions of the IgH locus in SETX^-/-^ cells which may be detrimental to class switching, contributing to the decreased class switching to IgA. Interestingly, GLT expression is often correlated with antibody isotype expression (77–79) and yet, we observed that SETX^-/-^ cells had difficulty class switching to the IgA isotype even with increased GLT expression in SETX^-/-^ B cells. This suggests that SETX^-/-^ B cells are receiving the appropriate upstream signals to induce class switching towards the IgA isotype and that the problem is likely downstream of GLT transcription.

Furthermore, B cells isolated from ATM deficient mice (ATM^-/-^) demonstrate elevated levels of microhomology usage, suggesting the use of a-EJ rather than NHEJ (80). Additionally, microhomology levels are drastically elevated in Sμ-Sɑ regions and are less evident at Sμ-Sɣ regions of B cells from A-T patients (80). The effect of SETX-deficiency on p-ATM foci induction following DNA damage may suggest that SETX-deficient mice are utilizing a-EJ rather than NHEJ during CSR, in a similar manner to ATM^-/-^ B cells. To explore whether SETX^-/-^ influenced the DNA repair pathways used during CSR, we sequenced the Sα region of cells that had switched to IgA using primers directed upstream of the Sμ region and downstream of the Sα region (37). While c-NHEJ is the default DNA repair pathway for S-region recombination (81) resulting in a microhomology of 0 to 4 bp (82), an alternative-EJ (a-EJ) pathway can also be used which relies upon longer microhomologies (83–85). Given that SETX interacts with 53BP1 in B cells (32), and 53BP1 is considered to promote c-NHEJ (86) we reasoned that a SETX deficiency may alter the repair pathway being utilized by SETX^-/-^ B cells. However, we observed no significant difference in DNA repair pathway choice between the SETX^-/-^ and SETX^+/+^ B cells. This may suggest the presence of other, redundant interactions in the NHEJ pathway that allow for its recruitment and usage even in the absence of SETX.

Herein, we have described impaired class switching to IgA in the SETX^-/-^ mouse model. These findings raise the possibility that individuals suffering from AOA2, who have loss-of-function mutations in SETX, may have a cryptic IgA-associated immunodeficiency. Environmental insults, including infection and the resulting inflammation, have long been thought to play a role in the onset and/or progression of neurodegenerative diseases (87, 88). It is possible that patients with AOA2 may experience elevated levels of inflammation early in life due to greater rates of infection of mucosal sites where IgA is highly enriched. Exploring this rare population of AOA2 patients, and their immune responses, merits further study and may shed light on AOA2 pathobiology.

## Acknowledgements

We thank Dr. Martin Lavin (The University of Queensland) for providing SETX^-/-^ mice. We would also like to thank Dr. Uttiya Basu (Columbia University) for providing the CH12-WT and CH12-SETX KO cells. This work was supported by an NSERC Discovery Grant and the Ontario Graduate Scholarship.

## Author Contributions

JM and VS conceived and performed the experiments. PZ performed computational analysis of immunoglobulin sequence diversity and switch region sequences. SA performed experiments and proof-read the manuscript. JA, JJCM, ATC, AM, JCA, KA, YT performed experiments. JM and VS wrote the manuscript in consultation with MRS and MSM.

## Conflict of Interest

The authors declare no competing interests.

---

**Supplemental Figure 1:**
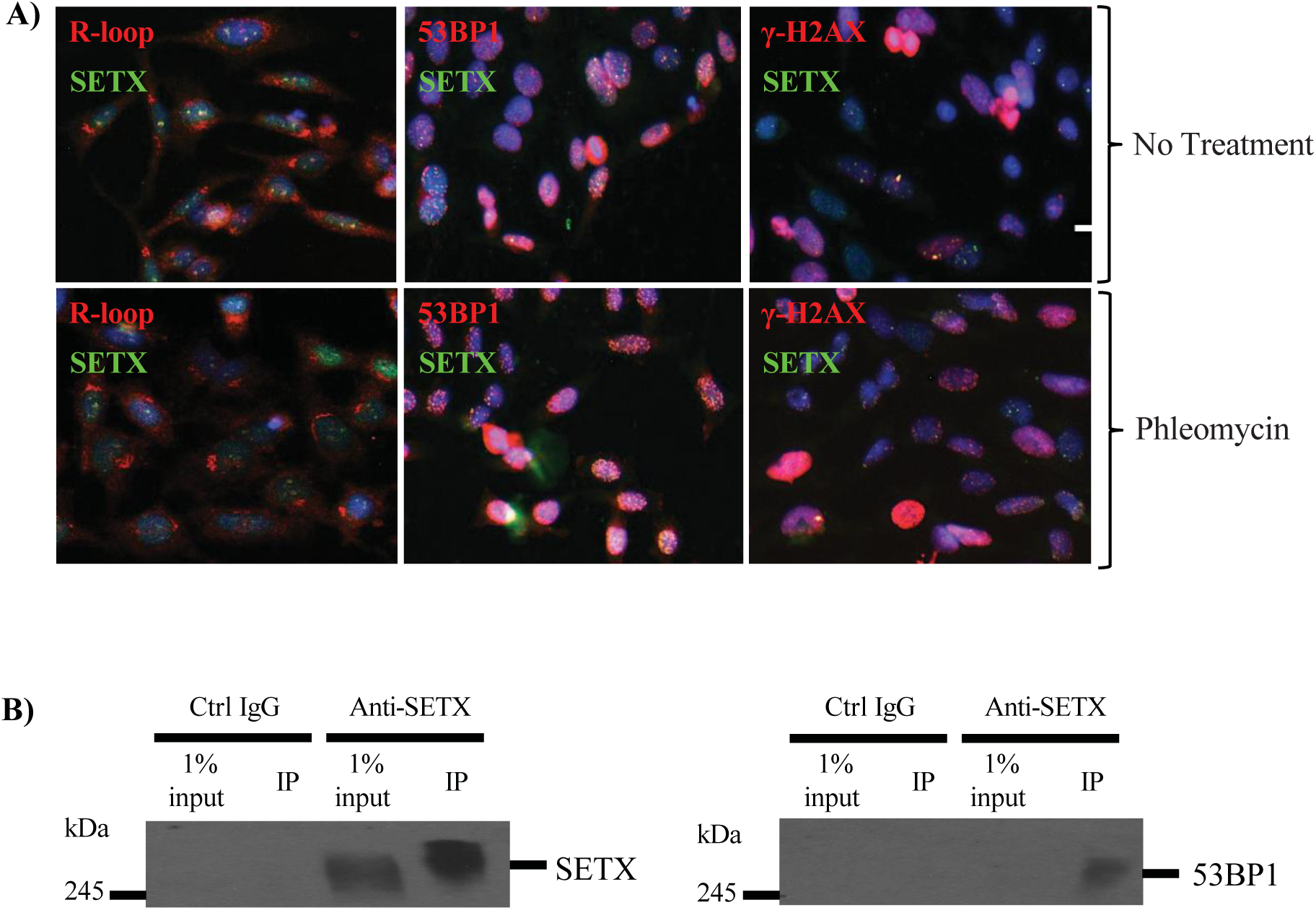
Senataxin interacts with common markers of the DDR. **(A)** HeLa cells were either untreated or treated with 20 µM Phleomycin for 1 hour and fixed with ice cold methanol. Cells were stained for SETX as well as either R-loops, 53BP1, or γ-H2AX. Cells were imaged with EVOS FL imager at 20X magnification. **(B)** Human PBMCs were isolated from peripheral blood using density centrifugation and stimulated for 3 days prior to protein isolation. SETX was immunoprecipitated and Western blot analysis was performed with anti-SETX and anti-53BP1.

**Supplemental Figure 2:**
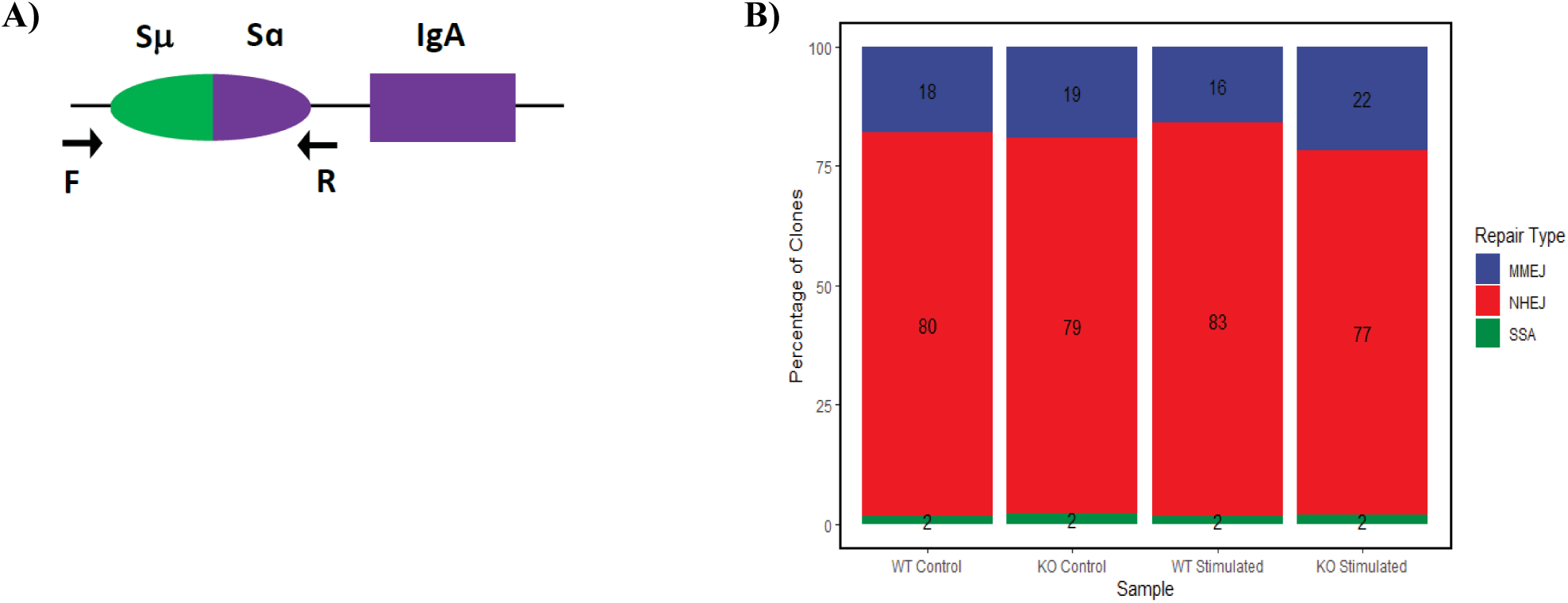
SETX does not alter the mechanism of the DDR during CSR. SETX^+/+^ (WT) and SETX^-/-^ (KO) CH12 cells were treated with IgA-specific stimulation (20 ng/mL IL-4, 1 ng/mL TGF-β, and 20 μg/mL LPS) or not treated (controls) for 5 days. **(A)** A graphical representation of the location of primers used for S-region sequencing. **(B)** DNA was isolated from WT and SETX KO CH12 cells after 5 days and sequenced using SMRT sequencing. DNA repair pathway choice was determined by quantifying sequence microhomology.

